# Proteome-wide comparison of tertiary protein structures reveal extensive molecular mimicry in *Plasmodium*-human interactions

**DOI:** 10.1101/2023.02.08.527763

**Authors:** Viraj Muthye, James D. Wasmuth

**Affiliations:** Faculty of Veterinary Medicine, University of Calgary, Calgary, Alberta, Canada; Host-Parasite Interactions Research Training Network, University of Calgary, Calgary, Alberta, Canada

**Author notes:** **Correspondence:** Corresponding Author: James D. Wasmuth.

**Keywords:** molecular mimicry, malaria, plasmodium, AlphaFold, tertiary structure, host-parasite interactions

## Abstract

Molecular mimicry is a strategy used by parasites to escape the host immune system and successfully transmit to a new host. To date, high-throughput examples of molecular mimicry have been limited to comparing protein sequences. However, with advances in the prediction of tertiary structural models, led by Deepmind’s AlphaFold, it is now possible to compare the tertiary structures of thousands of proteins from parasites and their hosts, to identify more subtle mimics. Here, we present the first proteome-level search for tertiary structure similarity between the proteins from *Plasmodium falciparum* and human. Of 206 *P. falciparum* proteins that have previously been proposed as mediators of *Plasmodium*-human interactions, we propose that seven evolved to molecularly mimic a human protein. By expanding the approach to all *P. falciparum* proteins, we identified a further 386 potential mimics, with 51 proteins corroborated by additional biological data. These findings demonstrate a valuable application of AlphaFold-derived tertiary structural models, and we discuss key considerations for its effective use in other host-parasite systems.

## Introduction

Parasites encounter host defenses at various points in their life cycle and employ a wide range of strategies for evading their host’s immune response and successfully transmitting to a new host (Chulanetra & Chaicumpa, 2021). These host-parasite interactions may be mediated by parasite-derived molecules—including proteins, lipids, sugars–that unexpectedly resemble host-derived molecules. This is termed ‘molecular mimicry’, which was originally defined as the sharing of antigens between parasite and host (Damian, 1964). One of the earliest reports of molecular mimicry came from the parasitic nematode *Ascaris lumbricoides*, which possesses A- and B-like blood group antigens in its polysaccharides (Oliver-González, 1944). The definition of molecular mimicry has adapted to keep up with molecular and genomic technologies and is now widely considered to similarity between proteins at the level of primary structure (amino acid sequence) and tertiary structure (summarized in (Tayal et al., 2022)). An assumption is that molecular mimicry confers a fitness benefit to the pathogen. However, in immunology research, the term molecular mimicry can be used to explain the cross-reactivity between exogenous and self-peptides and is the theoretical framework for understanding autoimmunity (Getts et al., 2013). Related to both these definitions, molecular mimicry might also result in heterologous immunity, in which the infection from one parasite protects against infection by other parasites with similar antigenic molecules (Balbin et al., 2023).

Here, our focus is molecular mimicry which likely confers a fitness advantage to the parasite, by either co-opting or disrupting the function of the mimicked host protein. Examples of molecular mimicry come from most branches of life. For instance, pathogenic bacterium *Escherichia coli* injects the TccP protein into host cells, which targets the polymerization of host actin. TccP contains multiple repeated motifs that mimic an internal regulatory element present in host N-WASP (neural Wiskott–Aldrich syndrome protein), which results in the activation of N-WASP (Sallee et al., 2008). This promotion of actin polymerization results in the creation of structures on epithelial cells that promote pathogen survival in the intestine. In another example, the myxoma virus decreases the number of activated macrophages by expressing its M128L protein on the host cell surface (Cameron et al., 2005). M128L shares significant sequence similarity with host CD47 and competes with it to bind with its receptor SIRPα. Within eukaryotic pathogens, the apicomplexan *Babesia microti* expresses the BmP53 protein which contains a domain that resembles thrombospondin (TSP1), a component of platelet cells (Mousa et al., 2017). The BmP53 TSP-1 is immunologically cross-reactive with human and it is proposed that BmP3 helps cloak the extra-cellular stages from the immune system.

To the best of our knowledge, the first study to identify host-parasite molecular mimicry at a genome-scale across multiple species was by Ludin and colleagues (Ludin et al., 2011). They considered the protein sequences from eight species of eukaryotic parasites, the host (human), and seven non-pathogenic, eukaryotic, negative control species. Their approach identified multiple potential instances of mimicry in these parasites. For example, they detected a 14 amino acid motif in multiple PfEMP1 proteins in *Plasmodium falciparum* that was identical to the heparin-binding domain in human vitronectin, a protein with multiple roles in human including cell-adhesion. The approach was repeated to find ninety-four potential mimicry proteins in a tapeworm-fish system (Hebert et al., 2015). It was also adapted and expanded for use with 62 pathogenic bacteria and identified approximately 100 potential mimics (Doxey & McConkey, 2013). These approaches rely on two proteins sharing enough sequence similarity to be detected by the sequence alignment software, *e.g*., BLAST (Altschul et al., 1990). However, proteins may share too little sequence similarity. For instance, several viruses express proteins with tertiary structure similarity, but undetectable sequence similarity to human Bcl-2, and interfere with regulation of apoptosis (Kvansakul et al., 2007; Westphal et al., 2007). Similarly, in *Plasmodium falciparum*, a search of parasite proteins targeted to host extracellular vesicles revealed that at least eight shared unexpected and significant tertiary structure similarity with host proteins (Armijos-Jaramillo et al., 2021).

The opportunity to detect host-parasite mimicry at the level of tertiary structure has been limited by the number of available tertiary protein structures. Even for a parasite as important as *P. falciparum*, the protein databank (PDB) contains structures from less than 4% of the protein-coding genes in its genome (Table 1). We expect that most, if not all, other bacterial and eukaryotic pathogen species will have worse coverage. Proteome-wide searches for host-parasite molecular mimicry at the level of tertiary structure depended on *in silico* predictions that were of inconsistent quality (Armijos-Jaramillo et al., 2021). The prediction of tertiary protein structures from amino acid sequences has seen a much-publicised boon, in no small part to the development of AlphaFold (Ronneberger et al., 2021). In an early large-scale application, the AlphaFold Protein Structure Database (AFdb) provided tertiary structure predictions for 16 model organisms and 32 pathogen species of global health concern (https://alphafold.com). Complementing the release of AlphaFold was Foldseek, a novel approach to aligning tertiary protein structures (van Kempen et al., 2022). Comparisons showed that FoldSeek was nearly 20,000 times faster than existing protein structure aligners while maintaining accuracy (but see (Holm, 2022)). These two major advances in structural bioinformatics—AlphaFold and Foldseek—have empowered us to investigate the usefulness of using tertiary protein structures for identifying instances of host-parasite molecular mimicry.

**Table 1.** The number of tertiary protein structures for each species used in this study. Structures were downloaded from two sources - the AlphaFold Protein structure Database (AFdb) and the RCSB Protein Data Bank (PDB). These structures were filtered according to the steps outlined in Section 2.1. The number protein-coding genes were taken from UniProt.

| Category | Species | Protein-coding genes in UniProt | AlphaFold structures (that passed filtering) | PDB chains | Uniprot proteins mapping to PDB structures | Total tertiary structures | Uniprot proteins with tertiary structures |
| --- | --- | --- | --- | --- | --- | --- | --- |
| Control | *Arabidopsis thaliana* | 27,498 | 20,460 | 4,560 | 899 | 25,020 | 20,612 |
| Control | *Caenorhabditis elegans* | 19,838 | 14,683 | 1,036 | 231 | 14,968 | 14,736 |
| Control | *Candida albicans*<br>SC5314 / ATCC<br>MYA-2876 | 6,759 | 4,777 | 285 | 62 | 5,813 | 4,790 |
| Control | *Danio rerio* | 26,355 | 19,541 | 1,026 | 157 | 20,567 | 19,627 |
| Control | *Dictyostelium*<br>*discoideum* | 12,727 | 8,259 | 345 | 55 | 8,604 | 8,266 |
| Control | *Drosophila*<br>*melanogaster* | 13,821 | 9,685 | 2,624 | 505 | 12,309 | 9,877 |
| Control | *Escherichia coli*<br>strain K12 | 4,402 | 4,220 | 11,242 | 1,087 | 15,462 | 4,332 |
| Control | *Glycine max* | 55,855 | 39,168 | 314 | 42 | 39,482 | 39,180 |
| Host | *Homo sapiens* | 20,594 | 17,226 | 125,073 | 8,215 | 142,299 | 17,568 |
| Control | *Methanocaldococ*<br>*cus jannaschii*<br>strain DSM 2661 | 1,787 | 1,708 | 511 | 99 | 2,219 | 1,709 |
| Control | *Mus musculus* | 21,968 | 16,911 | 13,419 | 2,227 | 30,330 | 17,562 |
| Control | *Oryza sativa*<br>*subsp. japonica* | 43,673 | 22,852 | 410 | 77 | 23,262 | 22,873 |
| Parasite | *Plasmodium*<br>*falciparum* isolate<br>3D7 | 5,372 | 2,909 | 1,417 | 207 | 4,326 | 2,949 |
| Control | *Rattus norvegicus* | 22,816 | 16,555 | 7,512 | 668 | 24,067 | 16,705 |
| Control | *Saccharomyces*<br>*cerevisiae* strain<br>ATCC 204508 /<br>S288c | 6,060 | 4,745 | 15,228 | 1,215 | 19,973 | 4,878 |
| Control | *Schizosaccharomy*<br>*ces pombe* strain<br>972 / ATCC<br>24843 | 5,122 | 4,224 | 1,388 | 295 | 5,612 | 4,260 |
| Control | *Zea mays* | 39,208 | 24,721 | 456 | 94 | 25,177 | 24,757 |

Our parasitic species of study is *Plasmodium falciparum*. While our understanding of the mediators of host-parasite interactions is limited, as the leading cause of severe malaria in humans, *P. falciparum* might represent the current pinnacle of our knowledge. Furthermore, the protein tertiary structures are complemented with a broad range of curated -omics datasets available on PlasmoDB, which can help with candidate prioritization (Amos et al., 2022; Aurrecoechea et al., 2008). Proteins expressed by *P. falciparum* mediate interactions with its human host at multiple stages in its life cycle (Acharya et al., 2017). Molecular mimicry plays a role at both the liver and blood stages for immune evasion and cytoadherence. For instance, RIFIN, a prominent erythrocyte surface protein expressed by *P. falciparum*, binds to human LILRB1 which inhibits stimulation of the immune response. RIFIN does this by mimicking MHC Class I, the activating ligand of LILRBA (Harrison et al., 2020). Meanwhile, the circumsporozoite protein (CSP), which promotes invasion of human liver cells, has an 18 amino acid region that is similar to a cytoadhesive region in mammalian thrombospondin (Cerami et al., 1992; Robson et al., 1988).

In this study, our goal was to identify *P. falciparum* proteins which share tertiary structure similarity with human proteins but not detectable sequence similarity. First, we examined *P. falciparum* proteins which are known or have been implicated to directly interact with human biomolecules. We found new potential instances of molecular mimicry. Second, we extended our approach to consider all *P. falciparum* proteins and leveraged experimental datasets to filter the candidate mimics. Overall, our study highlights the advantages of using tertiary protein structures for identifying instances of molecular mimicry.

## Methods and Materials

### 2.1. Compiling the datasets of tertiary protein structures

We downloaded tertiary protein structures for *Plasmodium falciparum* 3D7 (parasite), human (host), and 15 negative control species, *i.e*., species that are not infected by *P. falciparum* (Table 1, File S1). Protein structures for these species were downloaded from two sources - 1) the RCSB Protein Data Bank (PDB) (experimentally-determined protein structures, last accessed 06/16/2022), and 2) the AlphaFold Protein Structure Database (AFdb) (computationally-predicted protein structures, https://alphafold.ebi.ac.uk/). The structures downloaded from both sources were processed before analysis (explained below).

#### 2.1.1. Processing the PDB structures

Several PDB structures were composed of chains from multiple source organisms. We separated such structures into individual chains and extracted the appropriate chains corresponding to each species. For instance, the PDB structure 7F9N is composed of four chains (A to D), of which two chains (A and B) are from *P falciparum* Rifin (RIF, PF3D7_1000500) and two chains (C and D) are from the human leukocyte-associated immunoglobulin-like receptor 1 protein (LAIR1, Q6GTX8) (Figure 1A). We included only chains A and B for *P. falciparum*. Additionally, multiple structure chains were chimeric or ambiguous, *i.e*., mapping to multiple source organisms. For instance, the PDB structure 4O2X has two chains (A and B) which map to both *P. falciparum* and *Escherichia coli* strain K12. Such chains were discarded to not confound downstream analysis.

**Figure 1.**
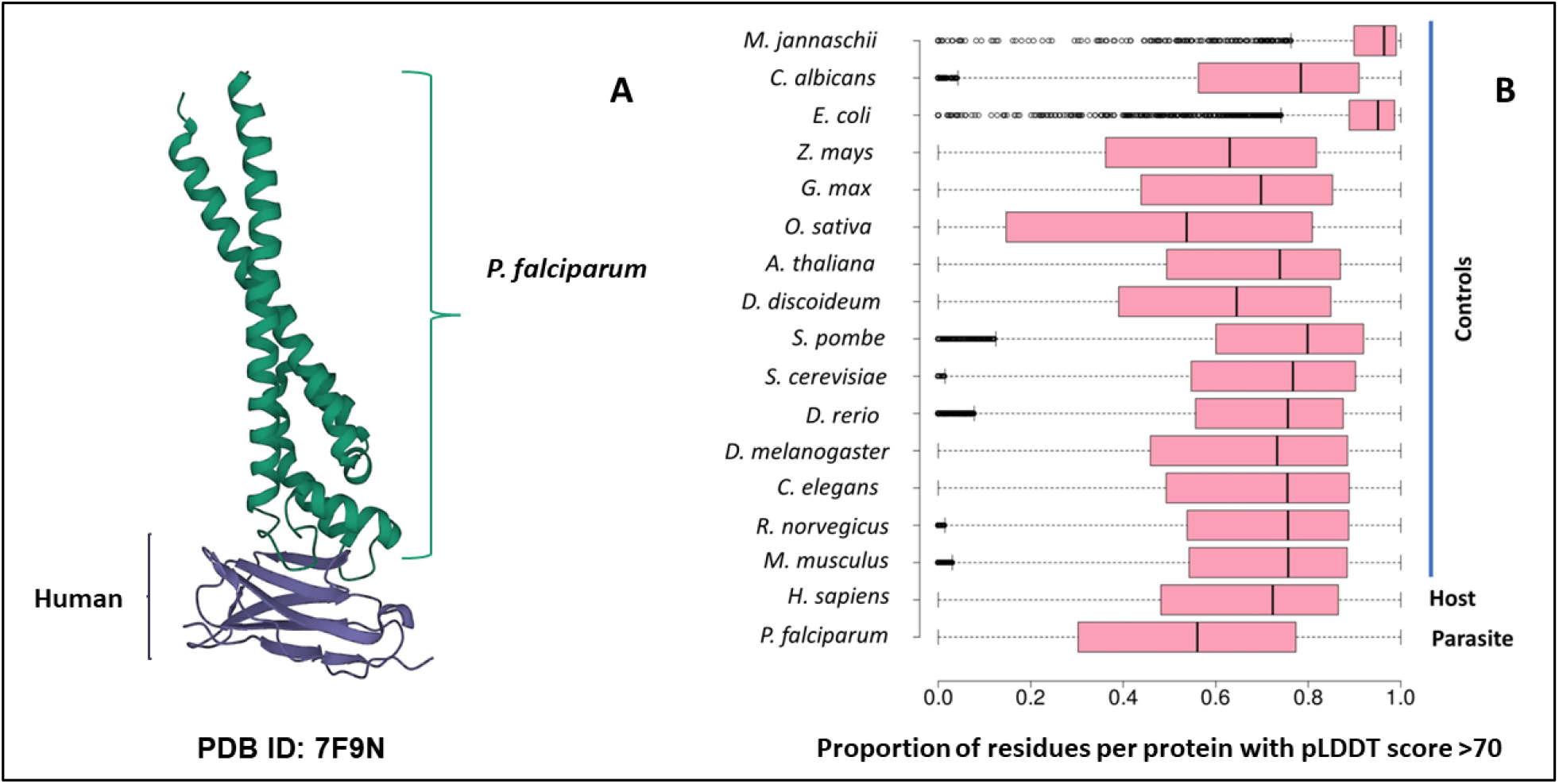
A) The 7F9N protein tertiary structure from the RCSB Protein Data Bank. The chains from the *P. falciparum* RIFIN protein (RIF, PF3D7_1000500) are colored in green and the chains from the human LAIR protein (LAIR1, Q6GTX8) are colored in purple. B) Box-plot representing the overall prediction accuracy for each protein in the AFdb. Each point in this distribution represents the proportion of all residues with a pLDDT score above 70 for one AlphaFold structure. The median value is the parallel bar within the box. The limits of the box are the 25^th^ and 75^th^ percentiles, the whiskers extend 1.5 times the interquartile range, and the dots are the outliers.

#### 2.1.2. Processing the AlphaFold structures

AlphaFold assigns a score to each residue in the predicted structure called the ‘pLDDT’ score. This score is a measure of prediction confidence for that residue. The pLDDT scores range from 0 (low confidence of prediction) to 100 (high confidence of prediction). Regions with a pLDDT score above 90 are modeled with high accuracy, between 70-90 are modeled well, and 50-70 are modeled with low confidence. The AlphaFold database suggests that regions with pLDDT scores less than 50 should not be interpreted as this low score could be indicative of intrinsic protein disorder. For each AlphaFold structure, we calculated the proportion of the total residues with a pLDDT score of more than 70. We retained predicted structures with at least half the residues of the structure modeled with a pLDDT score above 70 (Figure 1B, File S1).

### 2.2. Identification of *Plasmodium falciparum* proteins known to interact with human molecules

We performed a literature survey to identify *P. falciparum* proteins that are known to interact with human molecules. We started with a review of *P. falciparum*-human protein interactions (Acharya et al., 2017). Then, we identified all abstracts on PubMed using the query - [‘plasmodium falciparum’ AND ‘interact*’ AND ‘protein*’ AND ‘human*’] from 2017 to 2022. This resulted in 648 abstracts (as of 8/8/2022 1:36 PM). We read all 648 abstracts to identify the *P. falciparum* proteins of interest. The PlasmoDB ID for each protein was mapped to Uniprot IDs using PlasmoDB (Release 52, 30 August 2022). Three large gene families (PfEMP1, RIFIN, and STEVOR) play an important role in host-parasite interactions and pathogenesis in *P. falciparum*. Proteins belonging to these three families were downloaded from PlasmoDB.

### 2.3. Identification of sequence and structure similarity between *Plasmodium falciparum* and human proteins

#### 2.3.1. Analysis of protein sequence similarity

We analyzed sequence similarity between the proteins from *P. falciparum*, human, and 15 negative control species. We determined sequence similarity using three pairwise alignment search tools. SSEARCH36 implements the Smith-Waterman algorithm guaranteeing the optimal alignment. We used the following parameters for SSEARCH36 from Fasta36 - ‘m 8 -s BL62 -f 12 -g 1’. BLASTP (Altschul et al., 1990) and DIAMOND (Buchfink et al., 2021) implement heuristic algorithms that are faster than SSEARCH36 but do not guarantee the optimal alignment. We used BLASTP with an e-value cut-off of 1e^−3^ and performed an ultra-sensitive DIAMOND BLASTP search with the same e-value cut-off. For both aligners, we searched using the BLOSUM45 and BLOSUM62 substitution matrices. All other parameters were left as default. Additionally, we used OrthoFinder version 2.5.4 to identify groups of orthologous proteins between all the 17 species used in this study, using DIAMOND as the aligner (Emms & Kelly, 2019). The results of this analysis were used to identify human proteins that had orthologs in only the other three vertebrates used in this study (mouse, rat, and zebrafish).

#### 2.3.2. Analysis of protein structure similarity

We aligned all *P. falciparum* structures to a database consisting of human structures and control structures. The structural aligner used was Foldseek v4 (easy-search -s 9.5 --max-seqs 1000). We used an e-value cut-off of 0.01. Foldseek was also used to visualize structural alignments using the option ‘format-mode 3’.

#### 2.3.3. Expression analysis

Expression analysis of *P. falciparum* proteins was carried out using the publicly available RNA-seq datasets available in PlasmoDB. We identified all *P. falciparum* proteins with expression in the 90^th^ percentile in at least one of the stages in the intra-erythrocytic life cycle (young ring 8 hpi, late ring/early trophozoite 16 hpi, mid trophozoite 24 hpi, late trophozoite 32 hpi, early schizont 40 hpi, schizont 44 hpi, late schizont 48 hpi, and purified merozoites 0 hpi) using data from (Wichers et al., 2019). We also identified all *P. falciparum* proteins with expression in the 90^th^ percentile in the ring and/or sporozoite stage using data from (Zanghì et al., 2018).

## Results

### 3.1. Assembling and filtering the datasets of crystallised and computationally-predicted tertiary protein structures

We compiled tertiary protein structures from *Plasmodium falciparum* 3D7 (parasite), human (host), and 15 negative control species, those not infected by the parasite (Table 1, File S1). These structures were downloaded from the RCSB Protein Data Bank (PDB) and the AlphaFold Protein Structure database (AFdb). All AlphaFold-generated structures were filtered using the pLDDT score, a per-residue metric of the confidence of prediction accuracy. In line with the AlphaFold documentation, we considered structures to be high confidence if at least half their residues had a pLDDT score above 70. Through this filtering, we retained 56% of *P. falciparum* structures, 74% of human structures, and between 97% (*E. coli*) and 52% (*Oryza sativa*) for the control species (Table 1, Figure 1B, File S1).

### 3.2. Investigating the effect of the source of tertiary structures on Foldseek alignments

We wanted to determine whether the source of the tertiary structure—crystallised (PDB) or computationally-predicted (AlphaFold)—affected the Foldseek search results. Following our filtering steps, 167 *P. falciparum* proteins were represented by structures from both PDB and AlphaFold and 159 aligned to at least one structure from the host and/or negative control species. For each of these 159 proteins, we compared the Foldseek results for their PDB and AlphaFold structures. For most of these proteins (107/159), the results for both PDB and AlphaFold queries agreed between 90 and 100%. For almost 10% of these proteins (15/159), the agreement between the results was lesser than 50% (File S2, Figure S1).

### 3.3. Structural analysis of parasite proteins experimentally known to interact with human proteins

We performed a literature review and identified 74 *P. falciparum* proteins that interact with human molecules at various stages in its life-cycle (File S3). We also included three large *P. falciparum* gene families which are thought to play a role in parasite virulence—PfEMP1 (61 proteins), RIFIN (158 proteins), and STEVOR (32 proteins) (File S3). Overall, 206 of these proteins were represented by at least one structure in our database of PDB and high-quality AlphaFold structures. To understand whether molecular mimicry plays a role in how these proteins interact with the host, we asked the question: do these 206 *P. falciparum* proteins share sequence and/or tertiary structural similarity with human proteins?

We found that 31 proteins (15%) shared structural similarity with at least one human protein. Of these, three proteins aligned to human proteins which were restricted in vertebrates at the sequence-level (orthologs in only mouse, rat, and/or zebrafish). They were the parasite circumsporozoite protein (CSP, PF3D7_0304600) and two PfEMP1 proteins (PF3D7_0800100 and PF3D7_0617400). CSP was aligned to human thrombospondin (TSP1, P07996). Interestingly, as per previous sequence-based approaches, CSP mimics a cytoadhesive region in mammalian thrombospondin (Cerami et al., 1992; Robson et al., 1988).

Next, from these 31 proteins, we removed all parasite proteins which shared sequence similarity with human proteins – 15, 16, and 15 proteins aligned to human proteins by BLASTP, DIAMOND, and SSEARCH36 respectively (Figure 2A). Only 11 of these 31 *P. falciparum* proteins shared structure similarity but not sequence similarity to human proteins (Figure 2A). For these 11 proteins, we visually inspected their structural alignments with human proteins (File S4) and here we present the biological relevance of their interactions for seven proteins.

**Figure 2.**
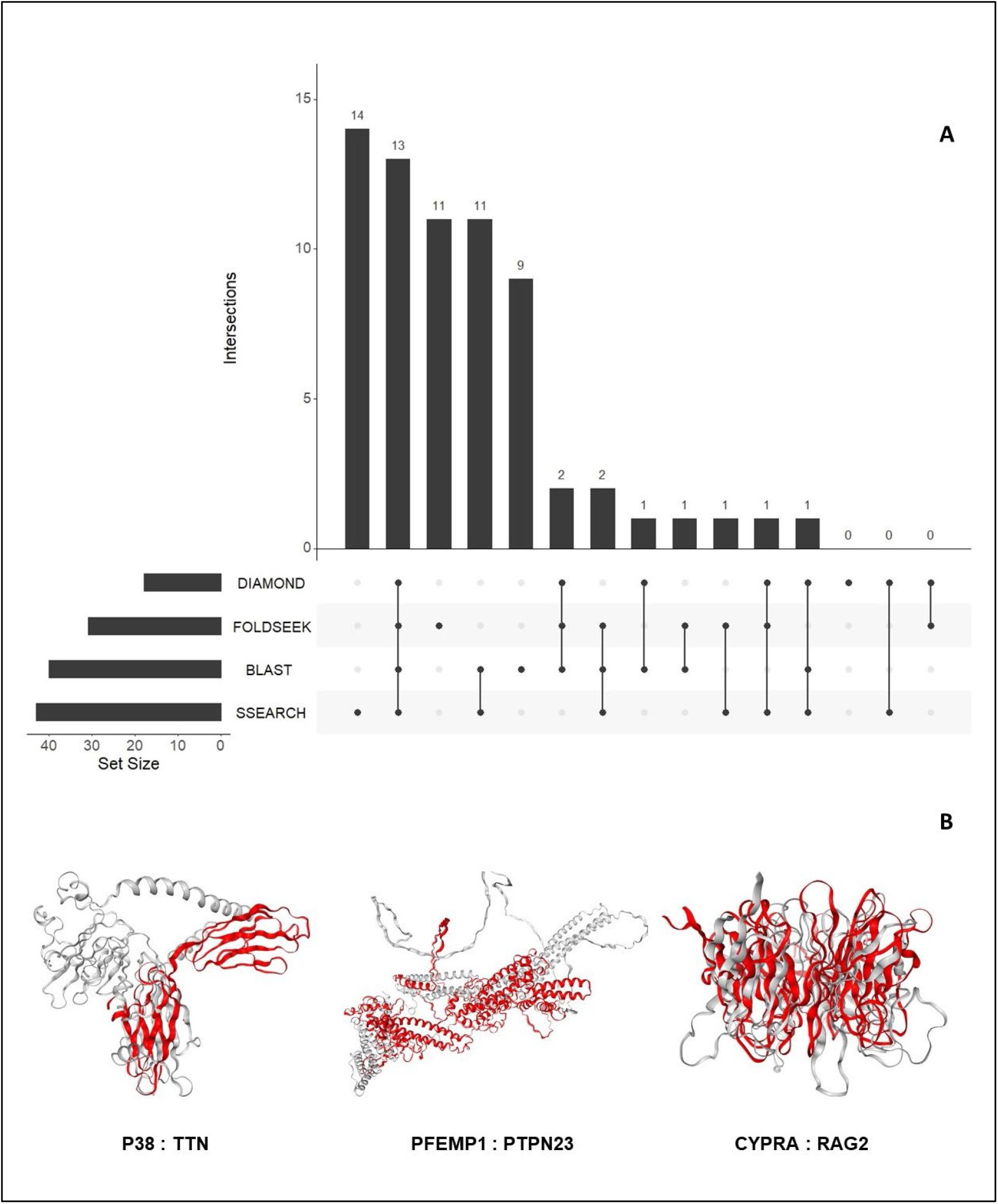
A) An UpSet plot of the 206 parasite proteins that interact with human molecules and are represented by at least one structure in our database. This plot displays the number of these interacting parasite proteins aligned to human proteins by DIAMOND, BLAST, Foldseek, and/or SSEARCH36. B) Tertiary structure alignments of the parasite structures (gray) and human structures (red). These alignments were generated and visualized using Foldseek.

**Figure 3.**
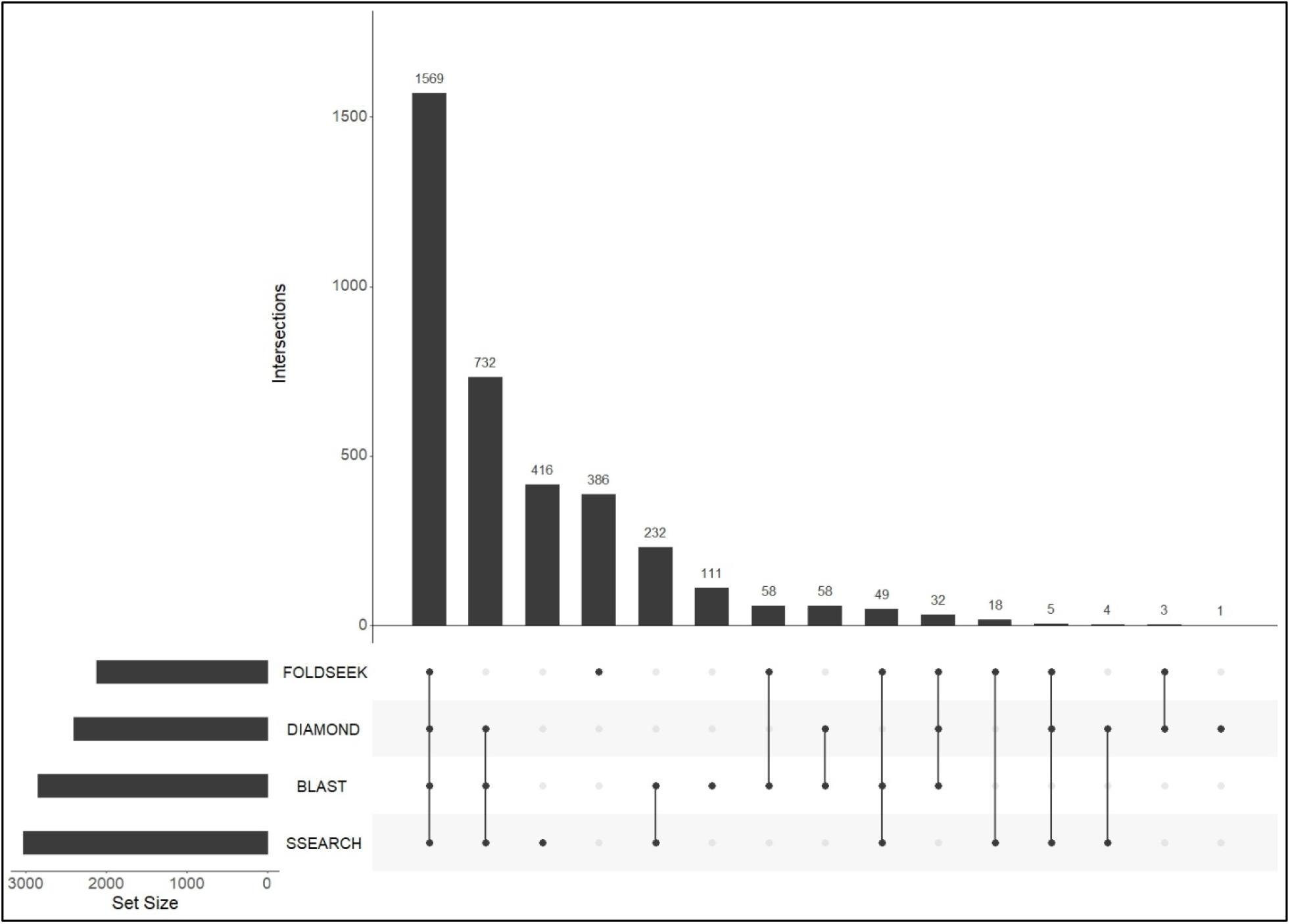
An UpSet plot of all *Plasmodium falciparum* proteins which were aligned to human proteins by Foldseek, DIAMOND, BLASTP, and/or SSEARCH36.

#### Three *PfEMP1* proteins

*P. falciparum* erythrocyte membrane protein 1 (PfEMP1) proteins primarily function in adhesion of infected erythrocytes to the vasculature. Here, our results suggest potential novel functions for three members of this large gene-family. Of the 61 PfEMP1 family proteins analyzed in this study, six shared structural similarity, but not sequence similarity, to human proteins. Two PfEMP1 proteins (PF3D7_0100300 and PF3D7_0600400) were similar to the human tyrosine-protein phosphatase non-receptor type 23 (PTPN23). Additionally, PTPN23 was the second-best Foldseek alignment for one more PfEMP1 protein (PF3D7_0937600). Interestingly, PTPN23 interacts with six human proteins that function in cytokine signaling (PTPN4, PTPN9, PTPN13, PTPN14, and GRAP2), suggesting that the three PfEMP1 proteins could be involved in immune modulation of the host by interfering in cytokine signaling (Figure S2).

#### AMA1

The apical membrane antigen (AMA1, PF3D7_1133400) interacts with the host erythrocyte membrane transport protein Kx protein (Kato et al., 2005) and is considered a potential vaccine target (Remarque et al., 2008). AMA1 has multiple crystallised structures, in addition to its AlphaFold prediction. AMA1 was aligned to human plasma kallikrein protein (KLKB1), a member of the plasma kallikrein–kinin system (KKS) in human. KKS proteins are known to interact with immune and complement systems (Wu, 2015). KLKB1 has also been implicated in the renin-angiotensin system (RAS), where it converts prorenin to renin (Schmaier, 2003). Some studies have reported an anti-malarial effect for RAS components. For instance, angiotensin II (which is formed by the cleavage of angiotensin to angiotensin I by renin and conversion to angiotensin II by angiotensin-converting enzyme) has been shown to inhibit different stages of multiple *Plasmodium* species and also reduce cerebral malaria pathogenesis (reviewed in (De et al., 2022)). We propose that AMA1 mimics KLKB1 to prevent the formation of angiotensin II by inhibiting renin activity.

#### P38

The 6-cysteine protein/merozoite surface protein P38 (P38, PF3D7_0508000) functions in red-blood cell invasion and binds to human glycophorin-A (GLPA, P02724) (Paul et al., 2017). We found that P38 was structurally similar to human titin (TTN, Q8WZ42) and fibroblast growth factor receptor 1 (FGFR1, P11362). The region from FGFR1 similar to P38 contained the two Ig-like protein domains ‘Immunoglobulin I-set domain (PF07679)’ and the ‘Immunoglobulin domain (PF00047)’. The regions similar to P38 from TTN also contained immunoglobulin-like domains. The ‘Immunoglobulin I-set domain’ are present in several adhesion proteins in human, where Ig-like domains play an important role in homophilic cell adhesion (Leshchyns’ka & Sytnyk, 2016). This suggests that P38, in addition to binding to GLPA, functions in adhesion owing to its similarity to the Ig-like domains.

#### CyPRA

The cysteine-rich protective antigen (CyPRA, PF3D7_0423800) interacts with two other parasite proteins (PfRH5 and PfRipr) to form a complex on the surface of an invading merozoite and is considered a potential vaccine target (Ragotte et al., 2020). CyPRA was represented by multiple structures from both the PDB and AFdb. The only human protein which shared tertiary structural similarity to both PDB and AlphaFold structures was the V(D)J recombination-activating protein 2 (RAG2, P55895) (Figure 2B). We identified the human interacting partners of RAG2 using StringDB (Figure S3). Seven interactors functioned in ‘signaling by Interleukins’, of which four were involved in ‘MAPK family signaling cascades’. The next best alignment was mitogen-activated protein kinase kinase kinase kinase 4 (MAP4K4, O95819), which also functions in the MAPK signaling. Thus, in addition to its key role in red blood cell invasion, structural similarity analysis suggests a possible novel role for CyPRA in modulating the host MAPK signaling pathway.

#### SHLP2

In at least one instance, our approach identified structural similarity in protein domains between the parasite and human proteins, which was missed by sequence similarity searches. Foldseek aligned the region containing the ‘Metallophos’ protein domain between the parasite protein ‘Shewanella-like protein phosphatase 2’ (SHLP2, PF3D7_1206000) and the human phosphatase ‘Serine/threonine-protein phosphatase with EF-hands 1’ (PPEF1, O14829).

### 3.4. Large-scale analysis of the structural similarity between *Plasmodium falciparum* and human proteins

The previous section shows that our approach can identify instances of molecular mimicry which mediate host-*Plasmodium* interactions. This motivated us to extend the approach to search all tertiary structures from *P. falciparum* against a database of 415,164 structures from human and the 15 negative control species. Of the parasite’s 4,326 tertiary structures, 3,649 (84%) were aligned to a structure from human and/or negative control species, and 3,350 (77%) could be aligned to a human structure.

Of these 3,350 structures, 59 structures were similar to only vertebrate proteins, and 27 aligned to only mammalian structures. Only eight structures aligned exclusively to a human structure. We further examined the structural alignments of these eight structures and identified potential novel biological functions for some of them. As mentioned above (section 3.3), the parasite protein CSP was aligned to human thrombospondin-1. The parasite protein 40S ribosomal protein S30 (RPS30, PF3D7_0219200) was shared tertiary structural similarity to human FAU ubiquitin-like and ribosomal protein S30 (FAU, P62861), which functions in apoptosis (Pickard et al., 2011) and is mapped to the GO term ‘innate immune response in mucosa’ in UniProt. This suggests that RPS30 could provide a biological advantage to *P. falciparum* by modulating the human immune response. The perforin-like protein 3 (PLP3, PF3D7_0923300) was aligned to human macrophage-expressed gene 1 (MPEG1, Q2M385), which functions in the host innate immune response. While PLP is expressed in both sexual and asexual stages of the parasite, it primarily functions in the mosquito-stage of the parasite (S. Garg et al., 2020). Thus, it is possible that PLP3 has an additional function in the human host to avoid the immune response.

Nearly one-fifth of the *P. falciparum* proteins that shared structural similarity to human proteins (386/2,120) had no detectable sequence similarity to a human protein (File S5). On average, 352 *P. falciparum* proteins had structural similarity but no sequence similarity to the control proteins. Overall, such *P. falciparum* proteins with structure similarity but not sequence similarity were not the same for each of these 16 species (Figure S4).

To prioritise the proteins, we categorized them with available annotations from PlasmoDB. Category 1 was predicted function, where the top scoring alignment was with a human protein with a GO term of interest—‘immune system process’, ‘cell adhesion’, ‘cytoskeleton’, or ‘signalling’. Category 2 was likely export from the cell, where the *P. falciparum* protein was predicted to contain a signal peptide. Category 3 was the gene’s expression, where we selected proteins whose genes was expressed in the 90^th^ percentile in at least one of the human life-cycle stages of the parasite—sporozoite and ring—or one of the intraerythrocytic stages (Wichers et al., 2019; Zanghì et al., 2018). A total of 169 *P. falciparum* proteins could be placed in at least one of the categories, with 97 proteins in category 1, 43 in category 2, 95 in category 3, 38 in two categories, and 13 proteins in all three categories (Figure S5, Table 2). Seven of the nine top human proteins aligned to these *P. falciparum* 13 proteins were taxonomically restricted to vertebrates.

**Table 2.**
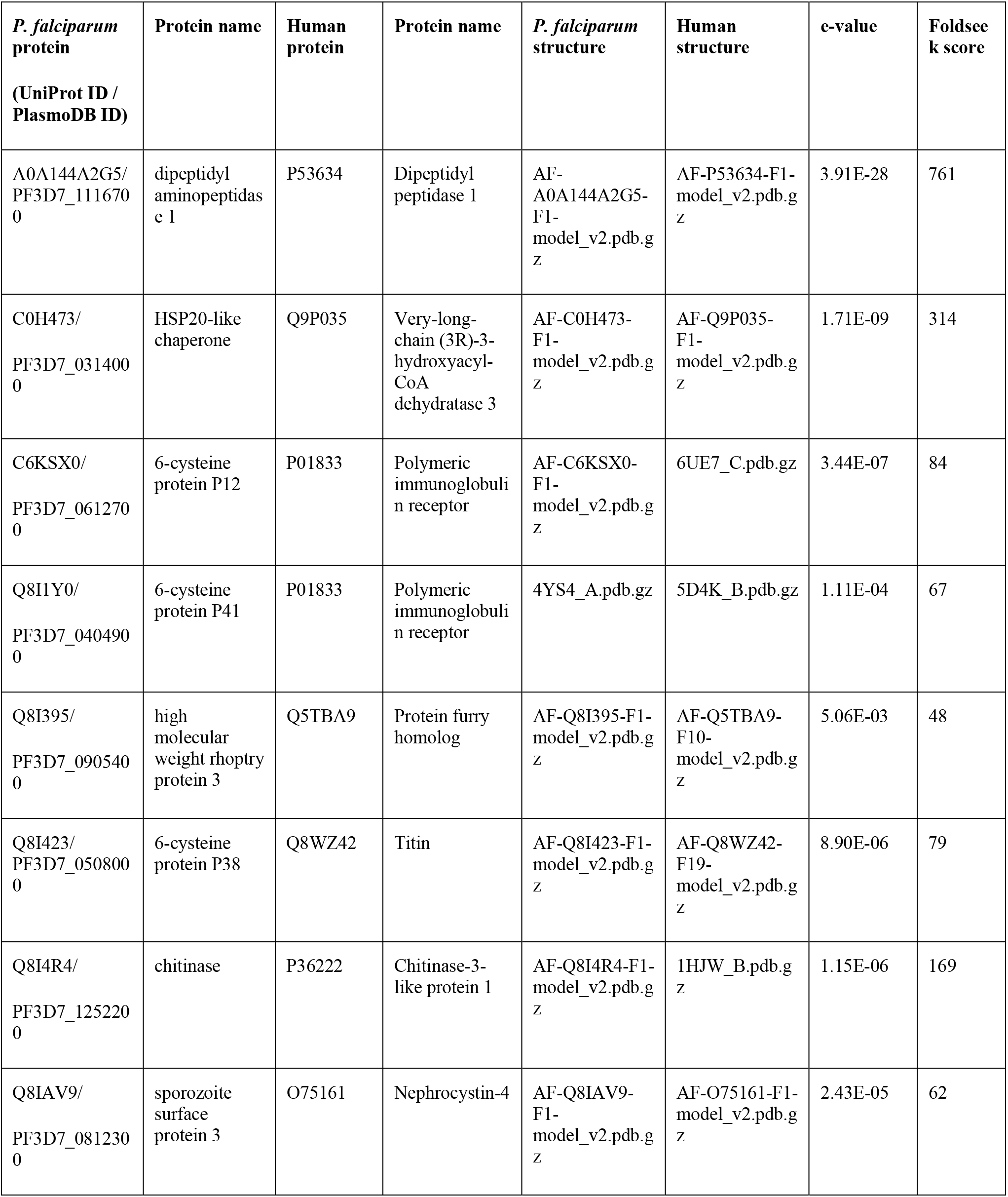
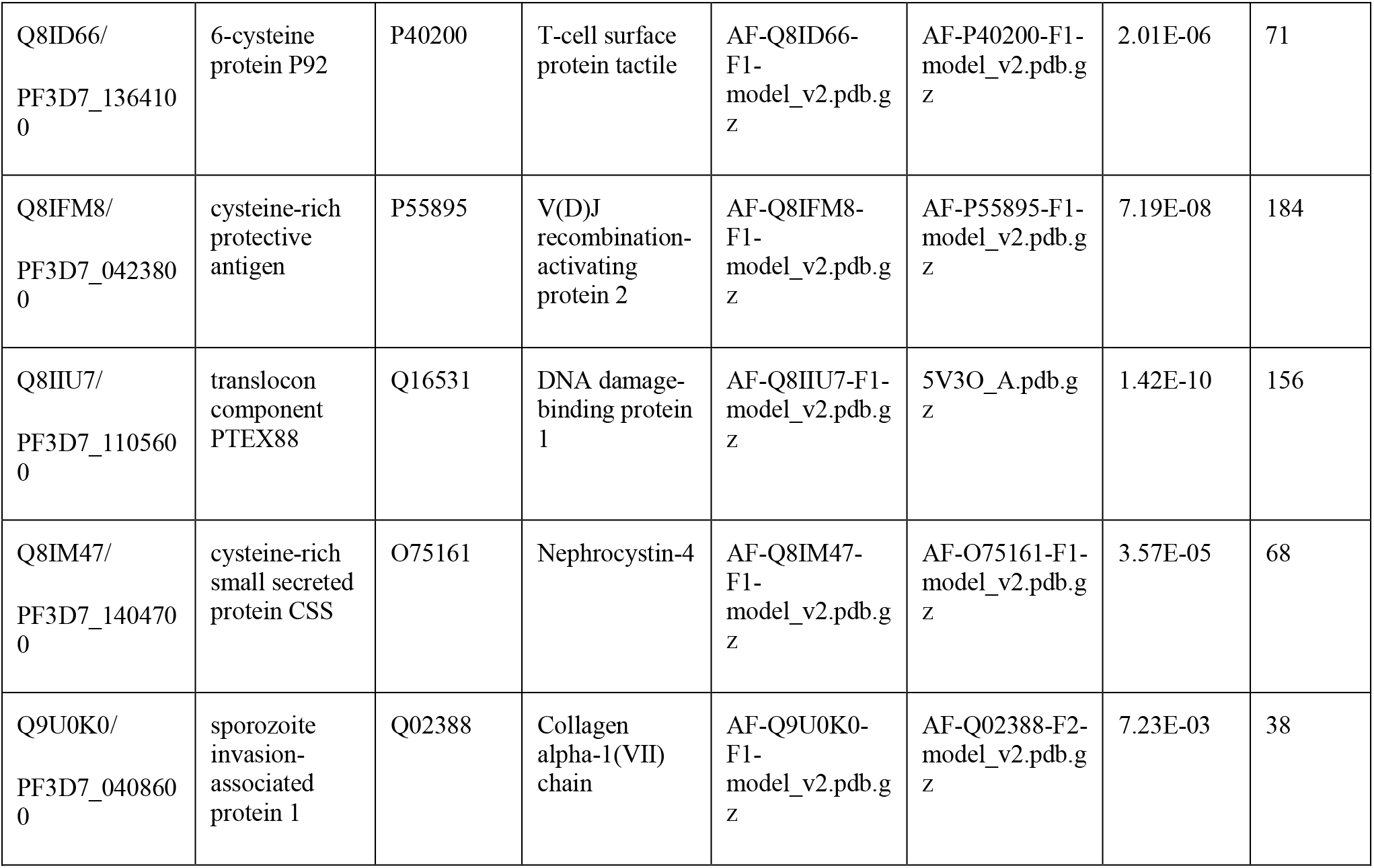
Top Foldseek alignments for the 13 P. falciparum proteins that were present in all three categories outlined in section 3.4.

The 51 *P. falciparum* proteins present in more than category represent instances of mimicry (File S6). Here, we present the biological relevance of a subset of these proteins. The four 6-cysteine proteins present in these 13 proteins are one of the most abundant surface antigens in the *P. falciparum* (Lyons et al., 2022) and have been attributed to virulence in related parasites (Wasmuth et al., 2012). These are promising candidates for mediators of host-parasite interactions. The first example was the P92/merozoite surface protein (P92, PF3D7_1364100), which interacts with human Factor H to downregulate the alternative complement pathway (Kennedy et al., 2016). We found that P92 shared structural similarity to human T-cell surface protein tactile (CD96, P40200), a transmembrane protein that is expressed by both T and NK cells (Georgiev et al., 2018). Interestingly, multiple roles for NK cells have been proposed at different stages of *P. falciparum* infection, including cytotoxic activity against the erythrocyte and hepatocyte stages (reviewed in (Wolf et al., 2017)). Additionally, six of the proteins CD96 interacts with function in immune response (GO:0006955) (Figure S6). Therefore, it is possible that P92 assists parasite infection by modulating the human immune response.

The second and third examples are the 6-cysteine protein P12/merozoite surface protein P12 (P12, PF3D7_0612700) and merozoite surface protein 41 (P41, PF3D7_0404900), which were similar to human polymeric immunoglobulin receptor PIGR (PIGR, P01833). PIGR is a known receptor for immunoglobulin M (IgM) (Pleass et al., 2016) and IgM is known to block the invasion of red blood cells by *Plasmodium* merozoites (Oyong et al., 2019). In fact, four other *P. falciparum proteins* - VAR2CSA, TM284VAR1, DBLMSP, and DBLMSP2, bind to IgM and affect IgM-mediated complement activation for evading the human immune response (Ji et al., 2022). Our structure-based approach suggests a similar role for P12 and P41. The fourth example is sporozoite invasion-associated protein 1 (SIAP1, PF3D7_0408600), which was aligned to human collagen alpha-1(VII) chain (COL7A1, Q02388). StringDB identified seven proteins that interact with COL7A1 and are mapped to the GO term ‘cell adhesion’ (GO:0007155) (Figure S7). Additionally, six proteins were mapped to the KEGG pathway ‘ECM-receptor interaction’ (hsa04512). Thus, we posit that SIAP1 could play a role in adhesion of the parasite to the host extracellular matrix.

An additional 35 parasite proteins were present in two of the three categories listed above. All warrant further investigation, but two are noteworthy. First, the sporozoite surface protein P36 (PF3D7_0404500) was similar to human PIGR, like two other members of the 6-cysteine protein family - P12 and P41. In fact, P36 was aligned to a similar region in PIGR compared to P12 and P41. This suggests that P12 potentially binds to IgM and helps the parasite evade the host immune response. When we analyzed all Foldseek alignments (not just the best hits), we found that an additional four 6-cysteine proteins were also similar to PIGR - 6-cysteine protein P12p (P12p, PF3D7_0612800), 6-cysteine protein P52 (P52, PF3D7_0404500), 6-cysteine protein (P48/45, PF3D7_1346700), and 6-cysteine protein P47 (P47, PF3D7_1346800). Second, the LamG domain-containing protein (PF3D7_0723200), which possesses a signal peptide, shared tertiary structure similarity to the human adhesion G-protein coupled receptor V1 (ADGRV1, Q8WXG9), a cell-surface protein involved in cell adhesion. StringDB found that five of its interacting partners function in ‘cell-cell adhesion’ (GO:0098609), suggesting that the *P. falciparum* protein assists parasite infection and survival by functioning in cell adhesion.

## Discussion

In this study, we present, to the best of our knowledge, the first genome-scale search of tertiary structural similarity between *P. falciparum* proteins and human proteins. We demonstrate the usefulness of our approach by showing that approximately 7% of the *P. falciparum* proteins had similarity to human proteins that could be detected only at the level of tertiary structure, and not primary sequence. Using available molecular and -omics datasets from PlasmoDB, we shortlisted a set of 51 instances of mimicry that are candidate mediators of host-parasite interactions.

We compared whether the source of the structural model made a difference. It made a considerable difference (<50% agreement) for 10% of proteins. We emphasize that this is for AlphaFold structures that were filtered for high confidence; we anticipate that for lower confidence AlphaFold structures, an agreement with a PDB structure would be worse. Improvement in the AlphaFold software and its evolutionary models will lead to better structural models, but users must remain vigilant of accuracy metrics when using this promising resource. If a particular gene of interest has a protein structural model available in the PDB, we recommend incorporating it in the analysis. A recent notable study attempted to overcome this challenge by improving AlphaFold predictions for two parasites - *Trypanosoma cruzi* and *Leishmania infantum* (Wheeler, 2021). They attributed the poor accuracy of tertiary structure prediction for several parasite proteins to the low number of representative parasite sequences used to predict them and demonstrated that they could improve the AlphaFold predictions by increasing the number of parasite sequences used to model these structures.

We validated our approach on known mediators of host-parasite interactions and we identified known similarity between CSP and thrombospondin-1. We also identified potential novel functions of multiple parasite proteins, which improves our understanding of existing *P. falciparum*-human interactions. One promising example involves multiple members of the 6-cysteine proteins found in *P. falciparum*. Members of this gene-family are expressed in multiple stages of the *P. falciparum* life-cycle in both the hosts (Lyons et al., 2022). Interestingly, while several members of this family have been implicated in critical functions like immune-evasion and host cell invasion, the precise roles for several of these proteins, like P12, P12p, P38, and P41, remain unknown (Lyons et al., 2022). Therefore, our structure-similarity-based approach improved our knowledge of how some of these proteins contribute to immune evasion, by identifying a potential binding-partner (IgM) and function for these proteins.

There are four important points to consider when interpreting the results of this analysis. First, we noticed that the number of parasite proteins with structural similarity, but not sequence similarity, to human and negative control proteins was similar for the majority of species studied. Some these would be distant homologs which are not detected by sequence-based tools employed in this study. Indeed, the failure of homology detection has been shown to be one of the important reasons for the presence of lineage-specific genes (Weisman et al., 2020). In fact, multiple studies have demonstrated the effectiveness of using structure data to identify distant homologs that are missed by sequence-based approaches (Andorf et al., 2022; Monzon et al., 2022). Other proteins, however, would represent false-positives of our approach based on our definition of ‘molecular mimics’, in which we define that a mimic confers benefit to the parasite. To this end, we used experimental data on *P. falciparum* from PDB to shortlist 46 proteins in *P. falciparum* that are structurally similar to human proteins and have a high probability of benefiting the parasite via its mimicry of a human protein. It is important to note that the majority of parasites lack similar resources, making such an analysis for them problematic. This highlights the need to develop resources, like PlasmoDB, for other parasites of global health concern.

Second, since we discarded AlphaFold predictions with low prediction accuracy, we ended up removing structures with a high level of intrinsic protein disorder. This is unfortunate as instances of mimicry have been identified in such disordered regions. For example, it has been proposed that viruses modulate host cellular processes by mimicking regions in disordered regions (Dolan et al., 2015; A. Garg et al., 2022; Xue et al., 2014). Such mimicking regions in disordered regions can be potentially identified by supplementing our structure-based approach with a *k*-mer based approach.

Third, the influence of the structure aligner must be considered. Foldseek is orders of magnitude faster than other currently available structural aligners and the only software that allows, in a reasonable time, the type of comparison presented here. However, the trade-off in terms of accuracy is a matter of debate (Holm, 2022; van Kempen et al., 2022). The alignment speed of DALI makes it unsuitable for large-scale structural similarity searches.

In conclusion, we present, to the best of our knowledge, the first genome-level search of tertiary structure similarity between *P. falciparum* and human proteins and use the results of the search to identify instances of molecular mimicry between the parasite and the host. The extensive experimental data for *P. falciparum* on PlasmoDB guided our efforts to unearth the biological relevance of the identified similarities. The list of *P. falciparum* mimics catalogued in our study represent excellent candidates for experimental validation. Our results help further our insights into the molecular nature of *Plasmodium*-human interactions and will be important to inform efforts on developing vaccines and therapeutics against malaria.

## Author Contributions

Conceived and designed the analyses: VM & JW. Performed the analyses: VM. Wrote the manuscript: VM & JW.

## Acknowledgments

We thank Kaylee Rich and Dr. Constance Finney for their discussions on various aspects of the manuscript. We acknowledge the high-performance computing resources made available by the Faculty of Veterinary Medicine and Research Computing at the University of Calgary.

## 12 Data Availability Statement

Supplementary materials for this study have been deposited in the Open Science Framework repository at Muthye, V., & Wasmuth, J. (2023, February 9). Data for Proteome-wide comparison of tertiary protein structures reveal extensive molecular mimicry in Plasmodium-human interactions.” https://doi.org/10.17605/OSF.IO/CUSYG

