## Supplementary Figures for "Proteome-wide comparison of tertiary protein structures reveal extensive molecular mimicry in *Plasmodium*-human interactions"

### 1 Supplementary Data

### 2 Supplementary Figures and Tables

#### 2.1 Supplementary Figures

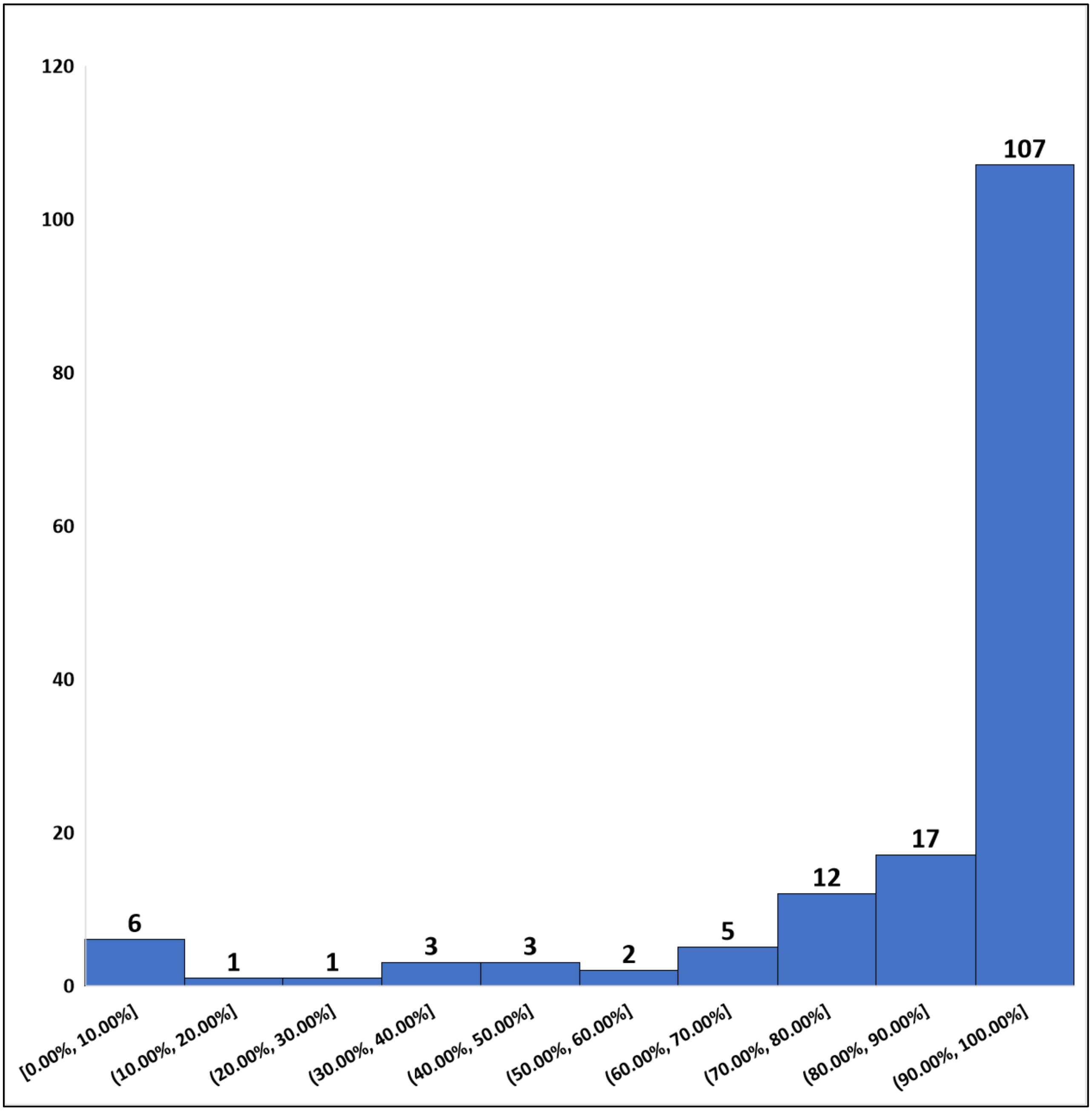


**Figure S1** Overlap between the number of (host and control) structures aligned to the AlphaFold structures and the number of (host and control) structures aligned to both PDB and AlphaFold structures.

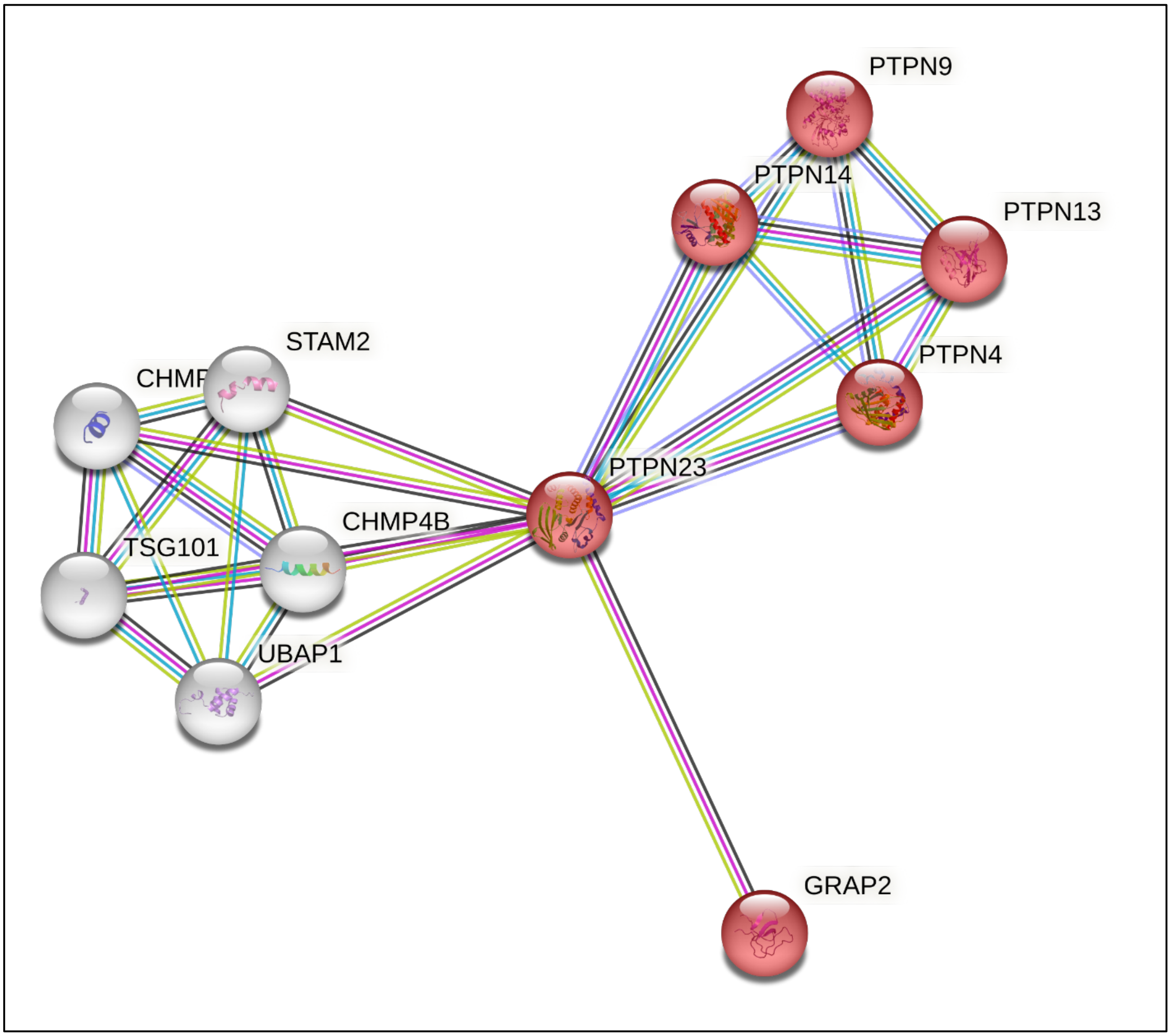


**Figure S2** StringDB analysis of the human tyrosine-protein phosphatase non-receptor type 23 (PTPN23). Red proteins function in 'Cytokine Signaling in Immune system' (Reactome pathway: HSA-1280215)

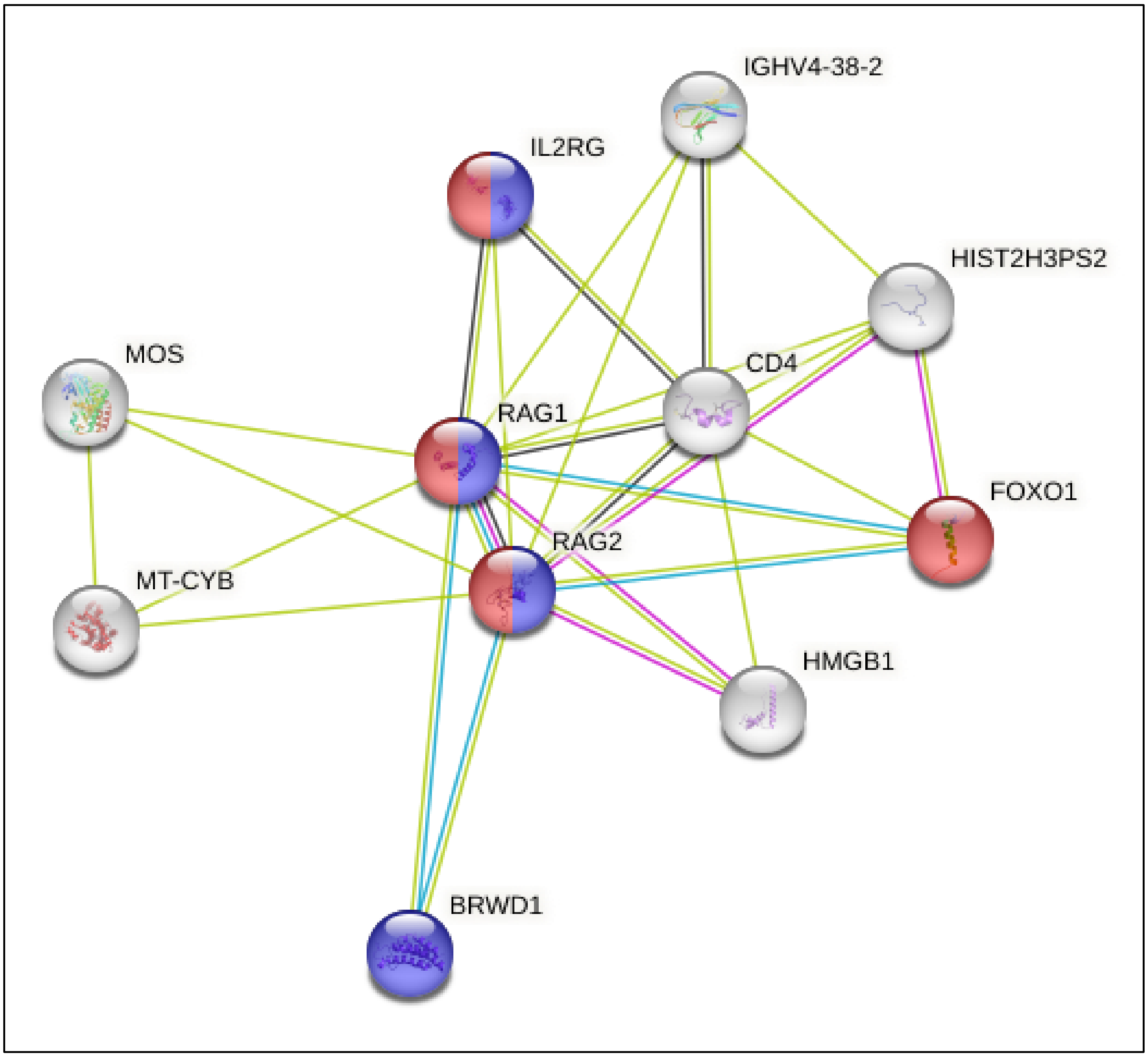


**Figure S3** StringDB analysis of the V(D)J recombination-activating protein 2 (RAG2, P55895). Red proteins function in 'MAPK family signaling cascades' (Reactome Pathway HSA-5683057) and blue proteins in 'Interleukin-7 signaling' (Reactome Pathway HSA-1266695)

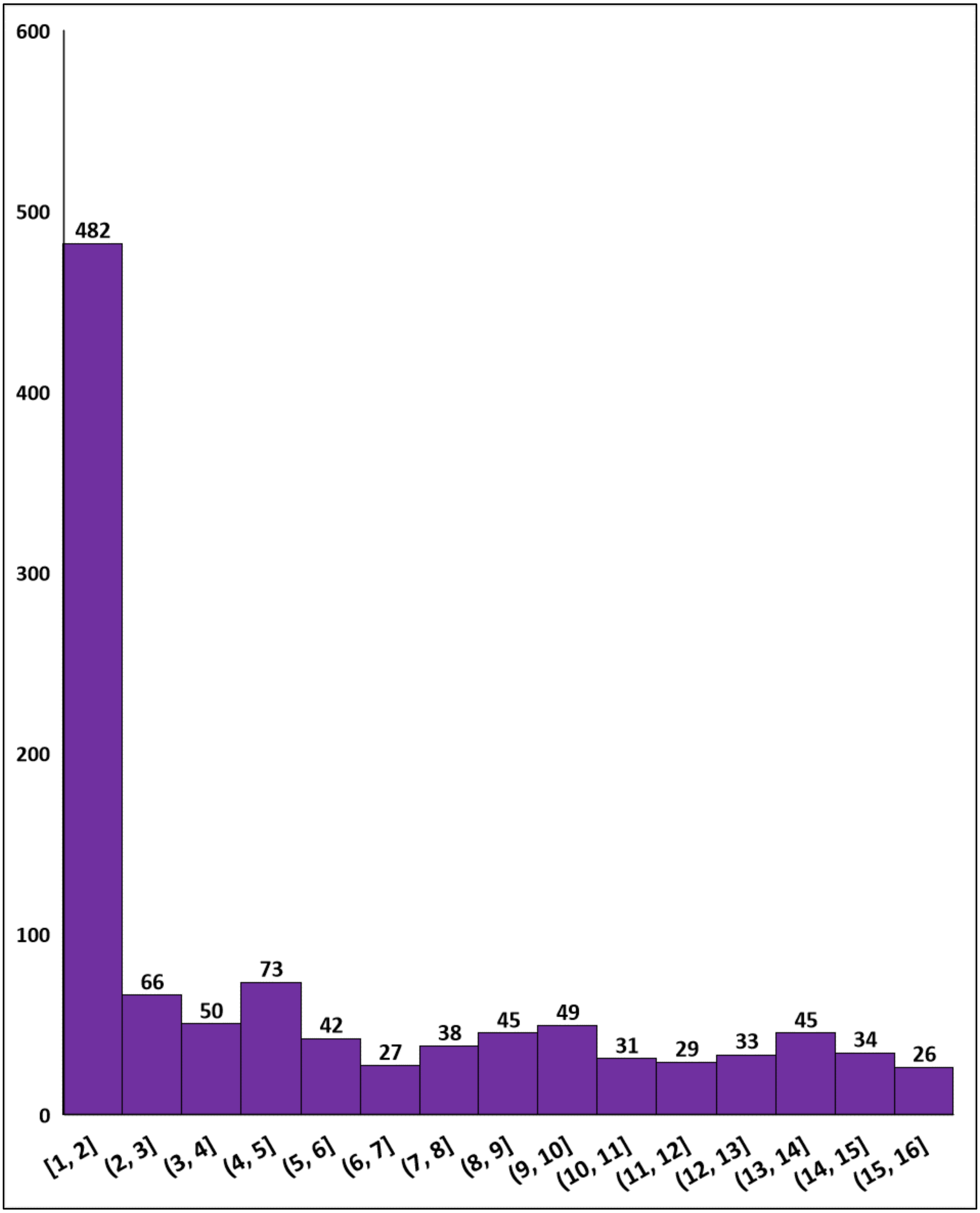


**Figure S4** For all 16 species, we identified parasite proteins which are aligned by Foldseek to at least one protein from that species, but not by BLAST, DIAMOND, or SSEARCH36. This is the distribution of the number of species each of those proteins was identified in.

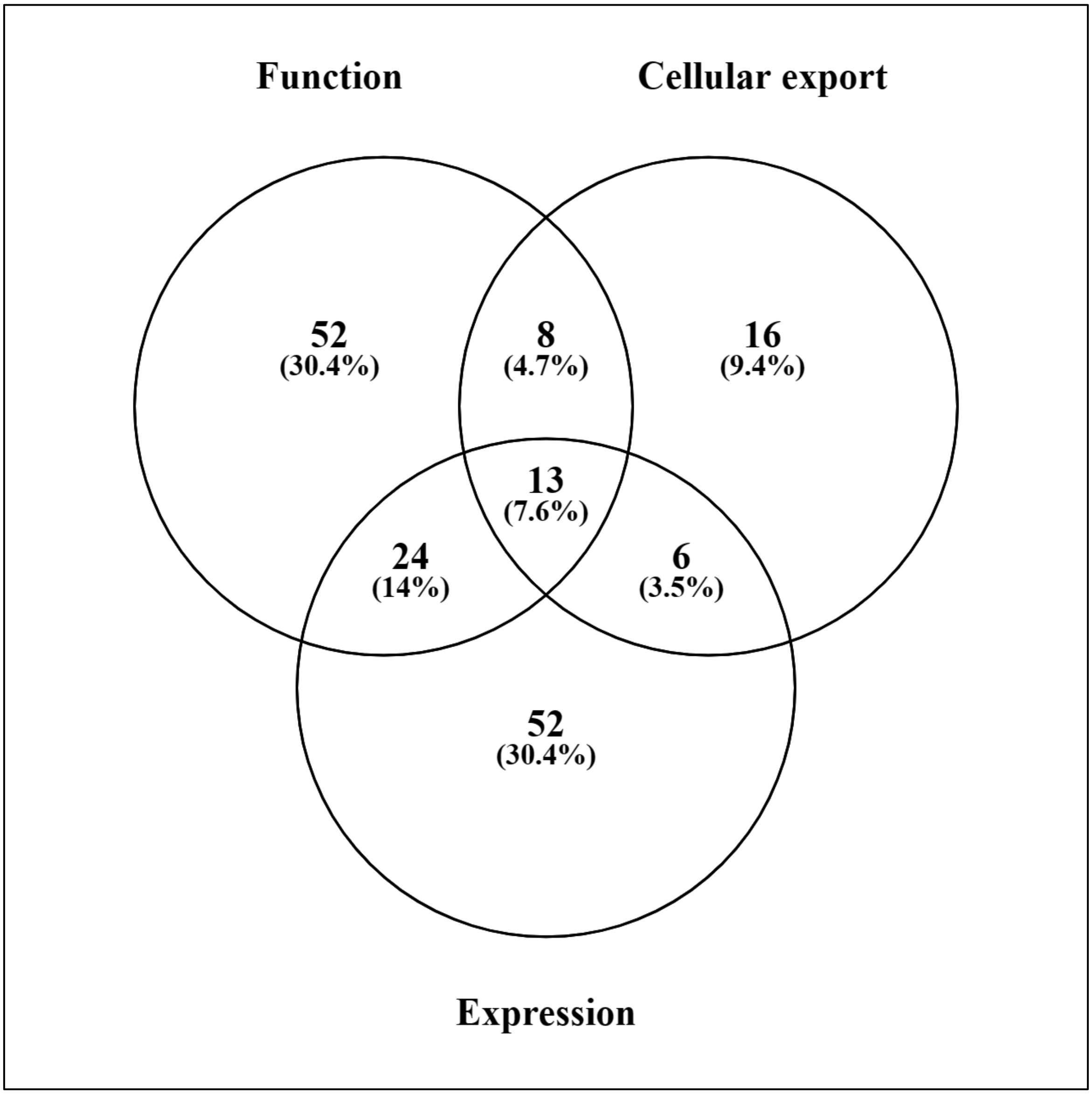


**Figure S5** Distribution of the number of parasite proteins identified by three categories listed in Section 3.4.

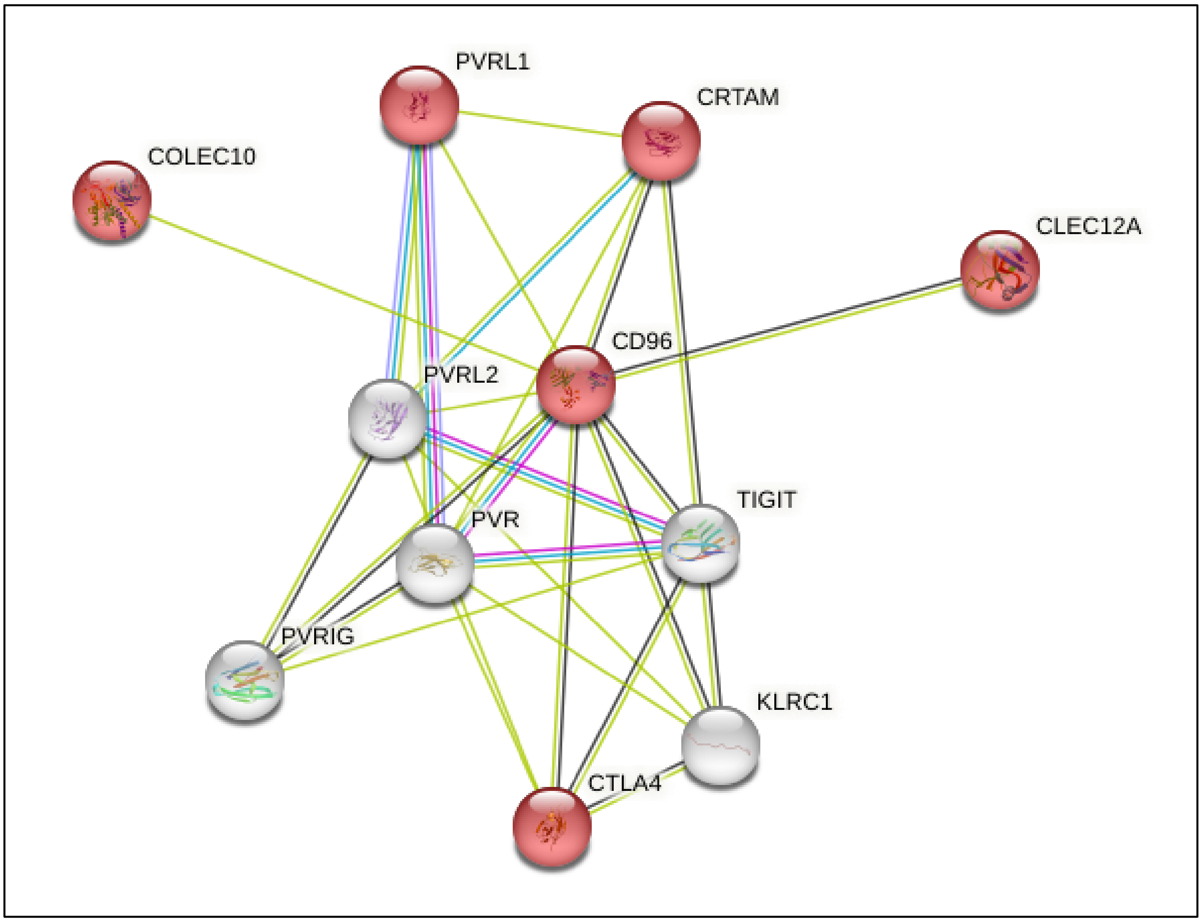


**Figure S6** StringDB analysis of the human T-cell surface protein tactile (CD96, P40200) where the proteins in red are mapped to the GO term 'immune response' (GO:0006955).

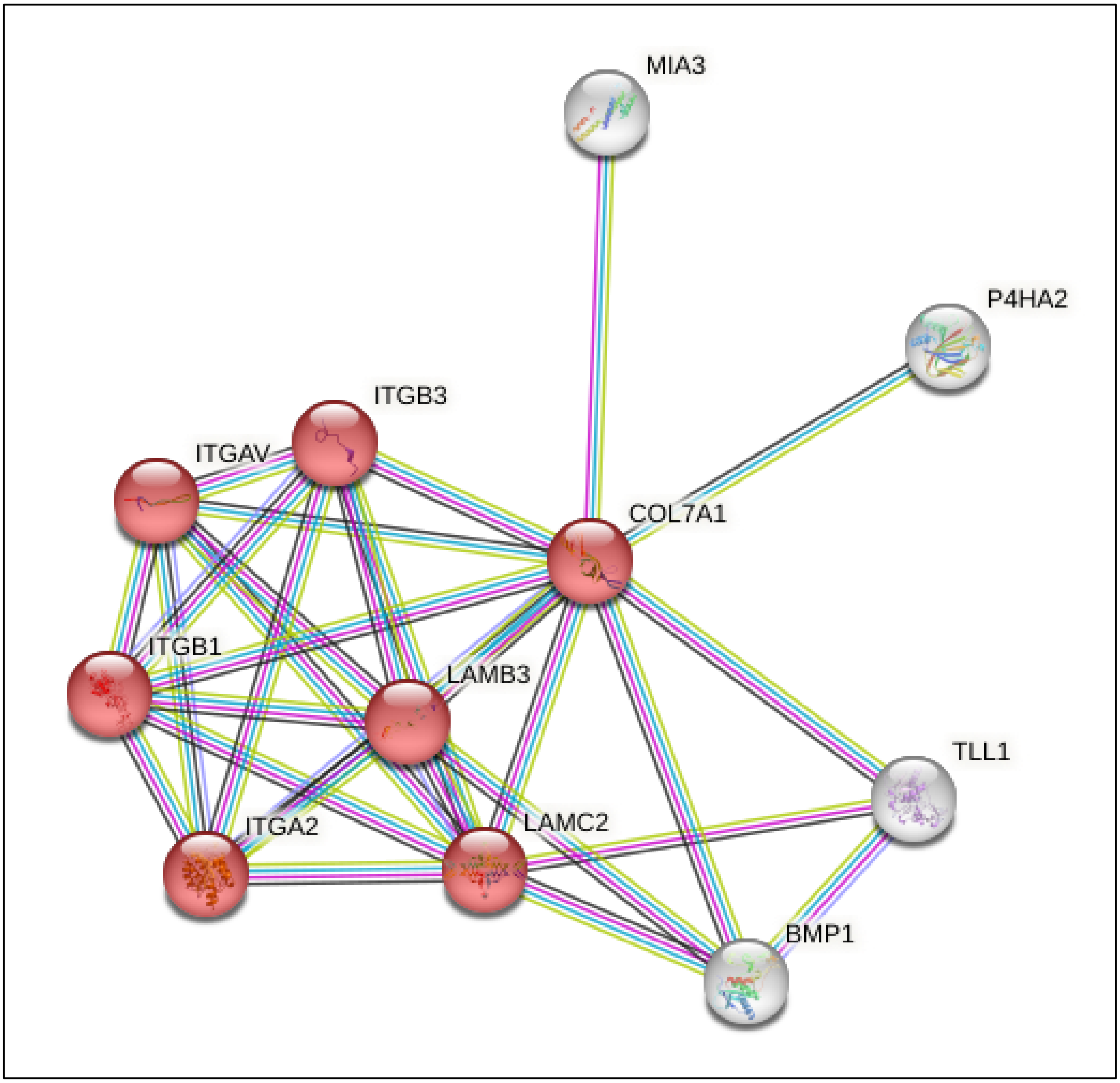


**Figure S7** StringDB analysis of the human collagen alpha-1(VII) chain (COL7A1, Q02388). where the proteins in red are mapped to the GO term ‘cell adhesion’ (GO:0007155).

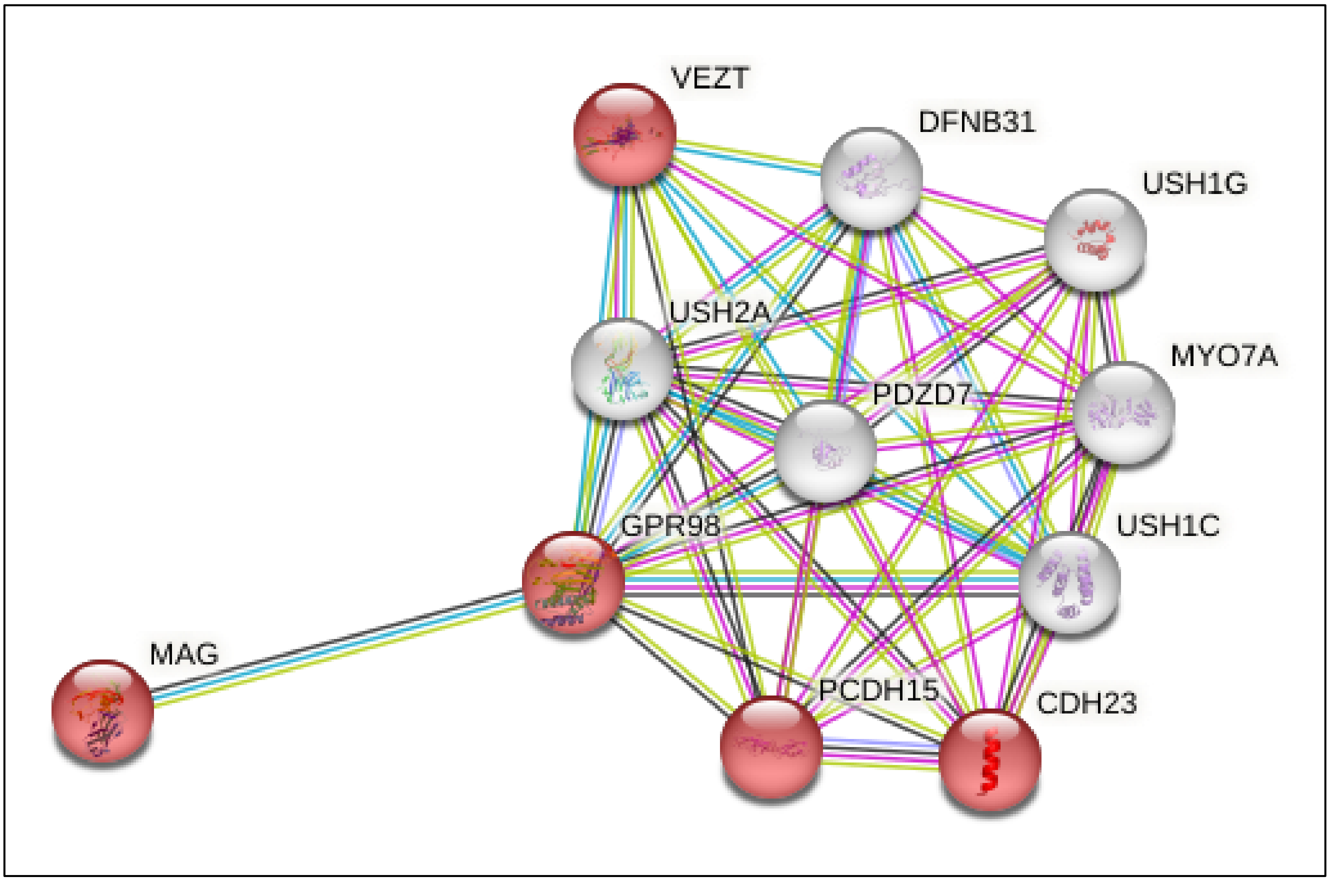


**Figure S8** StringDB analysis of the adhesion G-protein coupled receptor V1 (ADGRV1, Q8WXG9). where the proteins in red are mapped to the GO term 'cell-cell adhesion' (GO:0098609).
